# Correlation between plant cell wall stiffening and root extension arrest phenotype in the combined abiotic stress of Fe and Al

**DOI:** 10.1101/2023.06.20.545659

**Authors:** Harinderbir Kaur, Jean-Marie Teulon, Christian Godon, Thierry Desnos, Shu-wen W. Chen, Jean-Luc Pellequer

## Abstract

The plasticity and growth of plant cell walls (CWs) remain poorly understood at the molecular level. In this work, we used atomic force microscopy (AFM) to observe elastic responses of the root transition zone of 4-day-old *Arabidopsis thaliana* wild type and *almt1* mutant seedlings grown under Fe or Al stresses. The elastic parameters were deduced from force-distance measurements by AFM using the trimechanic-3PCS framework. In all metal stresses tested, the presence of single metal species Fe^2+^ or Al^3+^ at 10 µM exerts no noticeable effect on the root growth compared with the control conditions. On the contrary, a mix of both the metal ions produced a strong root extension arrest concomitant with significant increase of CW stiffness. This was not found for the *almt1* mutant which substantially abolishes the ability to exude malate. By raising the concentration of either Fe^2+^ or Al^3+^ to 20 µM, no root extension arrest was observed; nevertheless, a rise of root stiffness occurred. Our results indicate that the combination of Fe^2+^ and Al^3+^ with exuded malate is crucial for both CW stiffening and root extension arrest. However, stiffness increase induced by single Fe or Al metal is not sufficient for arresting root growth.

**Summary statement:** We record the change in stiffness of the external primary cell wall of living *Arabidopsis thaliana* seedlings in presence of metallic stress using atomic force microscopy. Results reveals for the first time the uncoupling between mechanical response (CW stiffening) and root extension arrest.

## Introduction

The cell wall (CW) of land plants has been depicted as a highly intertwining architecture by cellulose microfibrils, hemicellulose, and pectin (Carpita & Gibeaut, 1993), which compose the three major components of the primary CW. Cellulose microfibrils are the stiffest component, playing a load-bearing role, and their orientation creates a mechanical anisotropy, restricting cell expansion in the microfibril direction (Majda et al., 2017). Hemicelluloses (xyloglucan chains) bind to cellulose microfibrils using hydrogen bonds (Valent & Albersheim, 1974); they also bind covalently to pectin (Bauer, Talmadge, Keegstra, & Albersheim, 1973), a network made of matrix pectin polysaccharides and soluble proteins (Kerr & Bailey, 1934). Water is also a major constituent of primary CWs (Gaff & Carr, 1961), up to 65% (Jackman & Stanley, 1995), and an essential element for chemical reactions within the CW. The thickness of the primary CW was suggested around 80 to 100 nm for meristematic and parenchymatous cells, in accordance with a layered structure of cellulose microfibrils with a layer spacing of ∼20-40 nm (McCann, Wells, & Roberts, 1990). However, the accurate thickness measurement of external primary CWs remains challenging, roughly estimated as ∼0.1 to 1 µm (Derbyshire, Findlay, McCann, & Roberts, 2007).

Cell growth is characterized by an irreversible increase in cell volume and surface area, concomitant with a CW loosening. The complexity of CW growth results in poorly known pathways and mechanisms that control root CW plasticity (Somssich, Khan, & Persson, 2016). Upon various environmental stresses, a reduction of cell growth associated with CW stiffening is a well-known phenomenon observed in plants (Schopfer, 2006), tightly linked to dynamic behaviors of primary CWs. It has been proposed that strain-stiffening limits growth and restricts organs bulging (Kierzkowski et al., 2012). During the plant growth, some cells enlarge their volumes by 10-to-1000 times (Cosgrove, 1997) that is regulated by external stimuli such as temperature, light, water, xenobiotics, and internal factors like growth hormones (Preston & Hepton, 1960).

The cessation of coleoptile growth was attributed to the loss of CW plasticity but not to turgor pressure which implicates an increase of CW stiffness (Kutschera, 1996). One pioneer work on CW nanomechanics used atomic force microscopy (AFM) to observe stiffness heterogeneity in the meristem surfaces at regional, cellular and even subcellular levels (Milani et al., 2011). AFM has been shown powerful for stiffness measure on plant tissues (Cuadrado-Pedetti et al., 2021; Milani et al., 2014; Peaucelle et al., 2011). For characterizing the nano-stiffness of a sample in response to a given stress, AFM nanoindentation provides a promising strategy of detecting changes in physico-chemical properties of cellular or tissue surfaces on a nanoscale.

Recently, stiffening plant roots have been observed in the early 30 min after exposition to iron stress (Balzergue et al., 2017). In a condition of low phosphate, low pH (<6) and the presence of iron, a primary root extension arrest (REA) was observed and a signaling pathway involving STOP1 and ALMT1 proteins was found to inhibit CW expansions (Balzergue et al., 2017; Mora-Macias et al., 2017). Therein, STOP1 abundance in the nucleus of plant cell was found controlled by the presence of iron (Fe) and aluminum (Al) metals, of which both induced malate exudation through the ALMT1 channel (Godon et al., 2019; Le Poder et al., 2022). Although Fe is a fundamental nutrient for plants, a defense mechanism somehow occurs in a Fe-rich environment, implying that an excess of Fe is deleterious to plants (Oliveira de Araujo et al., 2020). The deleterious effect of Fe is linked with the ferritin capacity of plant cell for storing free reactive iron (Ravet et al., 2009) instead of being driven to the vacuole (Hirsch et al., 2006; Ward, Lahner, Yakubova, Salt, & Raghothama, 2008). Indeed, ferritin encapsulates the Fe^3+^ cation after oxidizing Fe^2+^ prior to storage (Macara, Hoy, & Harrison, 1972). In bean roots, the apoplast provides a storage space for Fe^3+^, where it could be extracted for nutrition use in case of iron deficiency (Bienfait, Vandenbriel, & Meslandmul, 1985). The *Arabidopsis lpr1/lpr2* mutants lack the capability of oxidizing Fe^2+^ to Fe^3+^ and were shown to reduce the amount of iron in the apoplast, exhibiting a Fe-insensitive phenotype in a low-phosphate condition (Svistoonoff et al., 2007).

Inhibition of root elongation is a well-known plant response to the tolerance of Al (Clarkson, 1965), especially at low pH (Bian, Zhou, Sun, & Li, 2013). Al toxicity resides in its cationic binding to negatively charged sites (membranes, proteins, saccharides) available in the root (Nichol, Oliveira, Glass, & Siddiqi, 1993). One creep-extension analysis showed that Al accumulation in the CW provoked a reduction of CW extensibility in wheat roots (Ma, Shen, Nagao, & Tanimoto, 2004).Within one hour of Al supply, callose deposition was observed in the root tip of soybean seedlings (Wissemeier, Diening, Hergenroder, Horst, & Mixwagner, 1992). In addition to callose deposition, the main physiological mechanism of Al tolerance is the exclusion of Al from the root apex (Kochian, Pineros, Liu, & Magalhaes, 2015), where Al usually accumulates in the root apex symplast and mostly in the apoplast (Delhaize & Ryan, 1995) and binds directly to negatively charged pectins of the CW of root border cells (Yang et al., 2016). This exclusion is accomplished by exudation of organic acids (Miyasaka, Buta, Howell, & Foy, 1991) such as malate and citrate (Liu, Magalhaes, Shaff, & Kochian, 2009). In cultured tobacco, Al accumulation in plant cell walls was found depending on the presence of ferrous iron (Fe^2+^) (Chang, Yamamoto, & Matsumoto, 1999). However, unlike Al, Fe does not stimulate malate excretion (Delhaize, Ryan, & Randall, 1993).

Fe^2+^ in phosphate-deficient conditions is able to arrest the primary root growth (Abel, 2011; Godon et al., 2019). Potential harmfulness of excessive Fe to cells is attributed to ROS (reactive oxygen species) production either by the Fenton (involving Fe^2+^) or by the Haber-Weiss reactions (Fe^3+^) (Gill & Tuteja, 2010). Above 40 µM of mixed Fe with Al resulted in a drastic reduction of root length, likely through the ROS production (Cakmak & Horst, 1991). The presence of Fe^2+^ in *Arabidopsis* roots stimulates ROS production with peroxidase activity (Balzergue et al., 2017; Muller et al., 2015; Naumann et al., 2022), particularly together with the class III peroxidase to stiffen and loosen the plant CWs (Francoz et al., 2015; Passardi, Cosio, Penel, & Dunand, 2005; Wolf & Hofte, 2014). In grass, peroxidase activity was linked to leaf growth arrest and CW cross-linking (MacAdam & Grabber, 2002); in rice, the peroxidase was found present in coleoptile growth arrest of shoots with increased ferulic and diferulic acids (Wakabayashi, Soga, & Hoson, 2012); similar findings were obtained for maize (Uddin et al., 2014). However, the causality between CW stiffness and REA remains to be elucidated.

In order to investigate Al and Fe effects on physiology and morphology of growing roots, we performed AFM indentations on *Arabidopsis* seedling roots under Fe and Al stresses of various metal concentrations and compositions. The present research provides a link of the structural stiffness measure (Chen, Teulon, Kaur, Godon, & Pellequer, 2023) with stress effects of metal ions in the root growth. Through the correspondence between the variations in the magnitude of elasticity parameters and the length of seedling roots in these stress conditions, it can improve our understanding of molecular mechanisms of metal ions in CW stiffening and root growth.

## Materials and Methods

### Seedling growth and manipulation

The experimental specimens are *A. thaliana* L. (Heynh.) lines of Columbia (Col-0) or the Col*^er105^* background as specified in (Bonnot et al., 2016). The production of *almt1^51^* mutant was previously described (Balzergue et al., 2017). Seeds were surface sterilized by 70% ethanol + 0.05% SDS for 1 min, followed by twice washing with 95% ethanol for 1 min each time and left in a laminar airflow for drying. To alleviate gravitropism effects on seedling growth such as inducing root wriggling or waving by growing vertically in a Petri dish, the seeds sown on day 0 were placed in a 24-well crystallographic plate (VDX plate HR3-140, Hampton Research). Plates were placed in a growing chamber (IPP100+ incubator, Memmert, Fisher Scientific, Illkirch, France) for 4 days with a 16-h photoperiod with 24°C/21°C day/night, respectively. During the 4 days, seedlings grew under the –Pi condition (no phosphate added) in the nutrient solution. The chemical content of the agar presently used is particularly poor in phosphate and metals, as determined by ICP analysis (Mercier et al., 2021), which is different from the agar used in our previous study (Balzergue et al., 2017).

After 4 days, seedlings were transferred into 60-mm agar Petri dishes in the –Pi condition while supplemented with or without 10 µM or 20 µM of FeCl2 and/or AlCl3for 2 hours. Then, seedlings were transferred from the agar plates to a glass slide for AFM nanoindendation experiments, and classical force–distance curves were collected within 30 min after mounting the glass slide. A control experiment, named No Transfer, in which seedlings were moved directly from the growing plate to the glass slide, was used to evaluate the impact of root transferring (from plates to Petri dishes).

### Length measurement of primary root

The root lengths were measured on day 6 after sowing with seedlings directly deposited in the Petri dishes. The photos were taken with a camera and the root lengths in the photos were measured using the NeuronJ plugin (Meijering et al., 2004) of ImageJ software (Schneider, Rasband, & Eliceiri, 2012) with a 5 mm grid paper as distance calibration. Snapshots from NeuronJ root tracing were saved in the PNG format and data were plotted using GraphPad Prism 5.0.

### Nanoindentation experiments with atomic force microscopy

Force–distance (F-D) curves were obtained using an AFM Dimension 3100 (Bruker, Santa Barbara, USA) with a nanoscope five controller running the Nanoscope 7.3 software. Triangular pyrex nitride cantilever with pyramidal tips of a max nominal radius 10 nm, a half-opening angle of 35°, and a nominal spring constant k = 0.08 N/m were used (PNP-TR, NanoWorld AG, Neuchatel, CH).

Calibration of photodiode sensitivity was done first using the approach-retract curve in air on the glass substrate followed by a thermal tuning to determine the cantilever spring constant (Kaur *et al*. 2023, submitted). The determined spring constants were about 0.08 ± 0.01 N/m. In case of a large divergence, the cantilever was manually readjusted inside the probe holder and the calibration was repeated (Schillers et al., 2017). Then the photodiode sensitivity was performed again in a liquid medium with an average value of 65 nm/V. In our case, a SUM value of 3.5-4 V was usually achieved with PNP-TR cantilevers. The engaging deflection setpoint was kept at 2.5 V while the initial vertical deflection on the photodiode was set to 0 V. For performing the indentation experiment, a ramp size of 3 µm, a scan rate of 0.5 Hz, and 4096 data points per curve were set. Trigger was set off and no trigger value was used, implying that z-start value for the ramp at each new engagement may need adjustments. To limit the maximal force during the measurement, a range of 25-40 nN was usually adopted for F-D data values.

The glass slide with a glued seedling (see the procedure in supplementary data and **Fig. S1**) was positioned under the AFM cantilever with the help of AFM optical camera. Thanks to the large motorized sample stage, we adjusted the glass slide in such a way that the cantilever could be positioned perpendicularly at the longitudinal middle of the glued root. The target working area, the transition zone, was 500 µm away from the root apex, almost twice the length of PNP cantilever.

### Hierarchical statistics and reproducibility of experiments

Owing to the roughness of root surfaces, indentations were performed at various locations in a matrix form. Different sizes of matrices were used: 5 × 2, 4 × 3, 4 × 4, representing 10, 12, or 16 indenting nodes. The optimal distance between nodes was 5 µm. Each node was formed of a sub-matrix with 3 × 3 or 2 × 2 F-D curves spaced by 50 nm in X and Y directions. Most of the presented results were obtained with a 4 × 4 node matrix of a 2 × 2 sub-matrix. Measurements from various forms of matrices were merged altogether. It usually took 25 min to record a full set of F-D curves for a single plant root; the manipulation time should be kept as short as possible to avoid additional stress effects.

For each stress condition, experiments were repeated 3 to 5 times. Each time involved 2 to 4 plants. The robustness of our protocol is ascertained by reproducible results from experiments repeated in a remote institute with another AFM instrument (Nanowizard IV, JPK-Bruker). Here, we considered all the measurements on one plant as one independent experiment. To synthesize the overall measurements into one comprehensive result for elasticity of the plant, hierarchical statistics were adopted. Explicitly, each elasticity parameter of one plant was obtained by averaging all the collected data (with 3 × 3 or 2 × 2 sub-matrices) of a node, then subsequently averaged over all the nodes of the plant. For one stress condition, at least 10 plants were analyzed (n ≥ 10).

Regarding the reproducibility of results, two criteria were imposed: a valid node should have more than half of its F-D curves within 2 sigma from the mean; a valid plant needs at least half of its measured nodes valid. The distribution of elastic parameters from all nodes of a given stress condition was most often log-normal. Therefore, we computed geometric means for the average value of elastic parameters of the plant. We also applied non-parametric Mann–Whitney t-tests to evaluating the statistical significance of these parameters among different stress conditions using a null hypothesis that assumes no difference on average among these conditions. A p-value was calculated using Graphpad Prism 5.0 with an α-threshold of 0.01. The box-and-whiskers plots were drawn using Graphpad Prism 8.

### Characterization of plant elasticity by the trimechanic-3PCS framework

The trimechanic-3PCS framework (Chen et al., 2023) allows us to investigate the variation of stiffness with varied depth for biomaterials of heterogeneous elasticity responding to an external force. For a depth of indentation trajectory exhibiting a linear-elasticity behavior, this theory states that the responding force *FT* of that depth zone can be expressed as a linear combination of three force components: *FC*, *FH* and *FS*. In this work, the elasticity parameters of the very surface of CWs, i.e., the first depth zone with depth *Z1*, are of concern.

The three force types (*FC*, *FH* and *FS)* govern three modes of restoration mechanics, namely, depth impact, Hookean and tip-shape nanomechanics, respectively. The contributions (or strengths) of the three nanomechanics to the overall response are represented by the spring constants (*k*C, *k*H, *k*S) of three parallel-connected spring (3PCS) analogs. The stiffness is defined as *k*T = *k*H + *k*S. Another important elastic parameter is *r*S = *k*S/*k*T, which quantifies penetration ease of the material and the composition of responding nanomechanics; it can represent material rigidity or deformability. Moreover, the *FS*–deduced effective Young’s modulus, *Ê* = E(1 − η^2^) with *E* the Young’s modulus and *η* the Poisson’s ratio, represents the intrinsic property of elasticity. The calculations of these parameters were detailedly described previously (Chen et al., 2023).

## Results

### Elasticity of WT seedling roots in the absence of metals

The non-stressed (control) systems were characterized as the seedling roots grown in no metals supplemented (Fe**0**Al**0**) with or without a transfer step. Systematically, the transfer step is always included, unless mentioned otherwise. Nanoindentation experiments were performed on these non-stressed seedlings 2 hours later after being transferred, or immediately in case of no transfer. The average of root length was obtained on day 6 (4 days growth + 2 days after transferring from the Fe**0**Al**0** condition) as 25.0 mm (**Table 1**). According to p-values with the significance threshold α = 0.01, no single group in the control systems significantly distinguishes itself from the others (**Fig. 1**). Averaged elasticity parameters are listed in **Table 2**. All pairs of groups tested by the standard non-parametric t-test were connected by a line with the exact p-value indicated above (**Fig. 1**). The results show that the averaged effective Young modulus *Ê* is about 55 kPa, the stiffness *k*T ∼4.64 mN/m, and indentation depth *Z1* ∼151 nm.

**Figure 1:**
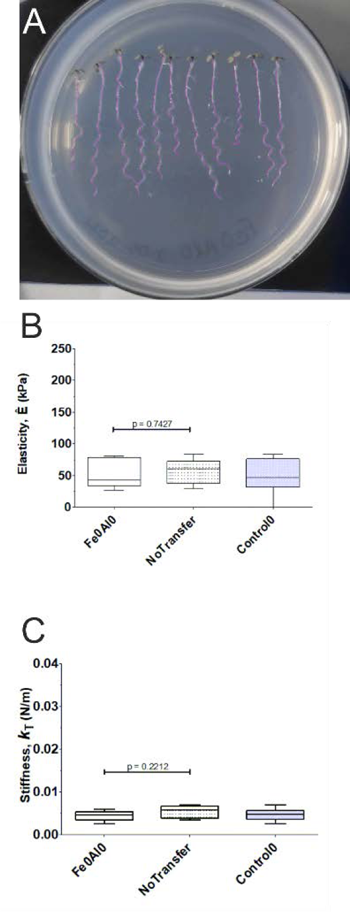
Phenotype and the box-and-whiskers plots of elastic parameters of WT seedling roots without metal stress. The whiskers represent minimal and maximal values, with the edge representing the first and third quartiles around the median, for each root system with stress condition specified. Fe**0**Al**0** indicates the nutrient solution for the system was not supplemented with Fe and Al. NoTransfer denotes seedlings that were not transferred from crystallography plates to Petri dishes, and elastic parameters were measured directly after being taken out of the growing plate. The control0 is referred to the overall results over both Fe**0**Al**0** and NoTransfer conditions. **A**) Snapshot of seedling roots, where length measurements were performed using the NeuronJ plugin of ImageJ. The purple color highlights the selected pixel used to calculate the root length. **B**) Effective Young’s modulus (Ê in the kPa unit) is presented for the control system. **C**) The values of stiffness *k*T in the N/m unit.

**Table 1:** The average of seedling root length for all study systems

| Plant type | Stress conditions | n | Length (mm) |
| --- | --- | --- | --- |
| WT | Fe**0**Al**0** | 19 | 25.0 ± 3.1 |
|  | Fe**0**Al**10** | 18 | 27.6 ± 3.1 |
|  | Fe**10**Al**0** | 17 | 26.8 ± 3.3 |
|  | Fe**10**Al**10** | 19 | 13.4 ± 1.3 |
|  | Fe**0**Al**20** | 26 | 26.8 ± 2.8 |
|  | Fe**20**Al**0** | 24 | 25.1 ± 2.2 |
|  | Fe**10**Al**10**+P | 27 | 25.0 ± 2.9 |
| *almt1* | ALMT1_Fe**0**Al**0** | 9 | 23.5 ± 2.1 |
|  | ALMT1_Fe**10**Al**10** | 9 | 22.0 ± 4.9 |

**Table 2:** Elastic properties of WT seedling roots

| | n | $Z_l$ (nm) | $\hat{E}$ (kPa) | $r_s$ | $k_T$ ($10^{-3}$ N/m) |
| --- | --- | --- | --- | --- | --- |
| Fe0Al0 | 11 | $147 \pm 55$ | $53.9 \pm 21.8$ | $0.78 \pm 0.05$ | $4.30 \pm 1.16$ |
| Fe0Al0_NoTransfer | 5 | $161 \pm 35$ | $56.3 \pm 20.1$ | $0.79 \pm 0.08$ | $5.40 \pm 1.45$ |
| Fe0Al10 | 10 | $150 \pm 62$ | $51.7 \pm 30.9$ | $0.74 \pm 0.07$ | $4.26 \pm 1.73$ |
| Fe10Al0 | 14 | $153 \pm 41$ | $58.4 \pm 50.4$ | $0.75 \pm 0.08$ | $4.94 \pm 3.51$ |
| Fe10Al10 | 11 | $127 \pm 57$ | $105 \pm 52$ | $0.81 \pm 0.09$ | $8.89 \pm 8.63$ |
| Fe0Al20 | 15 | $136 \pm 41$ | $76.9 \pm 39.4$ | $0.81 \pm 0.05$ | $5.72 \pm 2.18$ |
| Fe20Al0 | 11 | $119 \pm 29$ | $106 \pm 42$ | $0.83 \pm 0.06$ | $7.51 \pm 3.77$ |
| Fe10Al10+P | 8 | $162 \pm 25$ | $59.6 \pm 27.6$ | $0.81 \pm 0.06$ | $5.37 \pm 1.97$ |

## Elasticity of WT plant roots in the presence of metals

To assess the effect of the –Pi growth medium with Fe^2+^ and Al^3+^ ions, several combinatory concentrations of Fe and Al were applied to the test on the growth and CW stiffening of seedling roots. Systematically, the seedling roots were placed in various stressed environments for two hours. These stress conditions were prepared with 10 µM of FeCl2 or 10 µM of AlCl3, or mixing both, and labeled as Fe**10**Al**0**, Fe**0**Al**10** and Fe**10**Al**10**, respectively. No REA is observed in both Fe**0**Al**10** and Fe**10**Al**0** whereas a full REA is observed in the Fe**10**Al**10** condition (**Fig. 2A**), see T**able 1** for root lengths.

**Figure 2:**
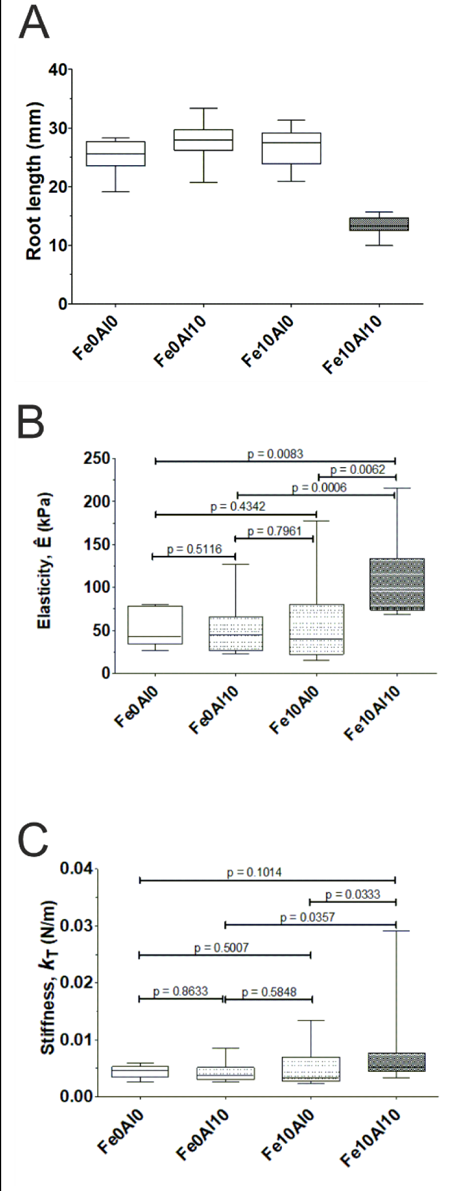
Box-and-whiskers plots of elastic parameters of WT seedling roots in the stress of 10 µM metal concentration. **A**) Average root lengths measured on day 6. **B**) Effective Young’s modulus (*Ê*) in the kPa unit. **C**) Stiffness measure, *k*T, in the N/m unit.

The results from nanoindentation experiments in 10 µM metal conditions are shown in **Fig. 2**. The hierarchical averages of their elasticity parameters are presented in **Table 2**. The elastic behaviors of conditions Fe**10**Al**0** and Fe**0**Al**10** exhibit no remarkable distinction from Fe**0**Al**0**; their combined result of *Ê* is about 55 kPa. It indicates that the total amount of metal ions at 10 µM changes little in the effective Young’s modulus, stiffness or indentation depth compared to no metal at all. However, the elasticity of roots grown with mixed Fe and Al (Fe**10**Al**10**) yields a value of 127 kPa for the effective Young’s modulus *Ê*, a significant increase in CW stiffness. Accordingly, the averaged *k*T of 8.89 mN/m for Fe**10**Al**10** is also much higher than all the conditions of a single metal element at 10 µM (cf. 4-5 mN/m in **Table 2**) (**Fig. 2C**).

We further explored the concentration impact of metal ions by doubling the concentration from 10 to 20 µM. We found that Fe**20**Al**0** displays a significantly higher *Ê* and *k*T than the control systems, while Fe**0**Al**20** exhibits a moderate effect (**Fig. 3**). The results show that the average of *Ê* and *k*T have a similar value between Fe**20**Al**0** and Fe**10**Al**10** conditions (**Table 2**) whereas the corresponding values of Fe**0**Al**20** are intermediate. Very interestingly, doubling the cationic concentration of single metal does not provoke the occurrence of REA (**Table 1**).

**Figure 3:**
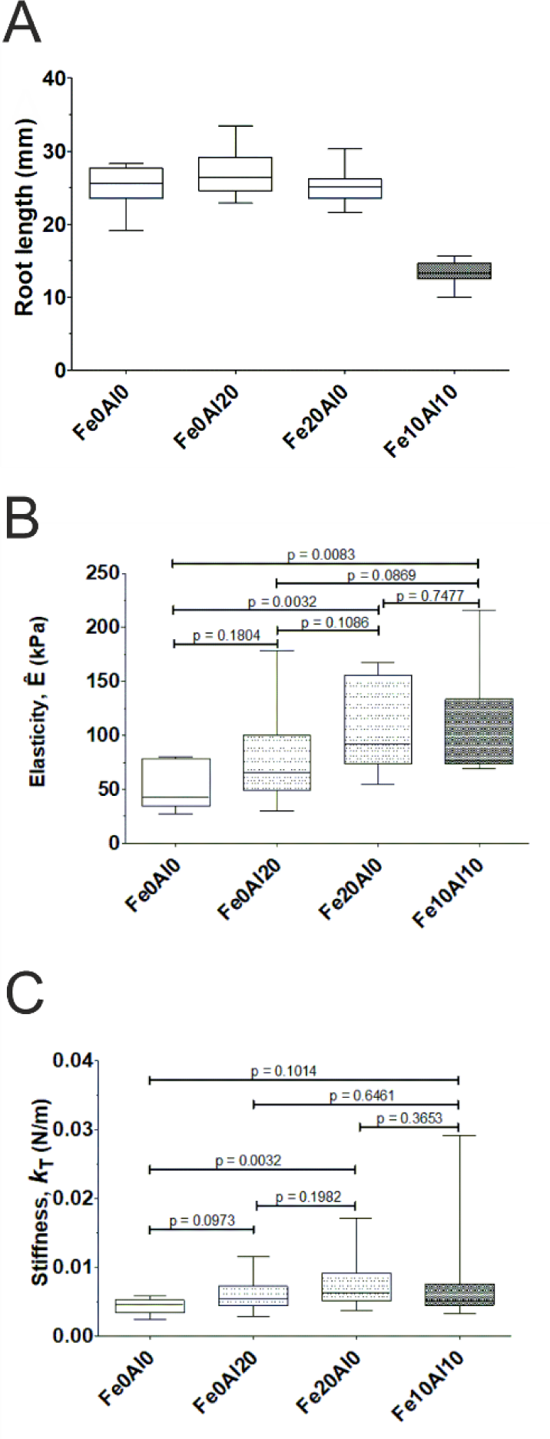
Box-and-whiskers plots of elastic properties of WT seedling roots with 20 µM of metallic ions. For comparisons, the results from WT Fe**0**Al**0** and Fe**10**Al**10** are co-presented. **A**) Average root lengths measured on day 6. **B**) Effective Young’s modulus (*Ê*) in kPa. **C**) Stiffness *k*T in N/m.

### Elasticity of *almt1* mutant plant roots in the presence of metals

Unlike WT plant roots, no REA phenotype was found from *almt1* mutants in the Fe**10**Al**10** condition (**Fig. 4**). The elasticity parameters for *almt1* mutant seedlings grown in Fe**0**Al**0** and Fe**10**Al**10** conditions are listed in **Table 3**. No significant difference was found in the magnitudes of *Ê* and *k*T between the two stress conditions (**Fig. 4**). However, these values are comparable to WT in Fe**0**Al**0** (**Tables 2, 3**). It implies that without exuded malate, the two metal ions cannot exert substantial effects on elastic responses of mutant roots.

**Figure 4:**
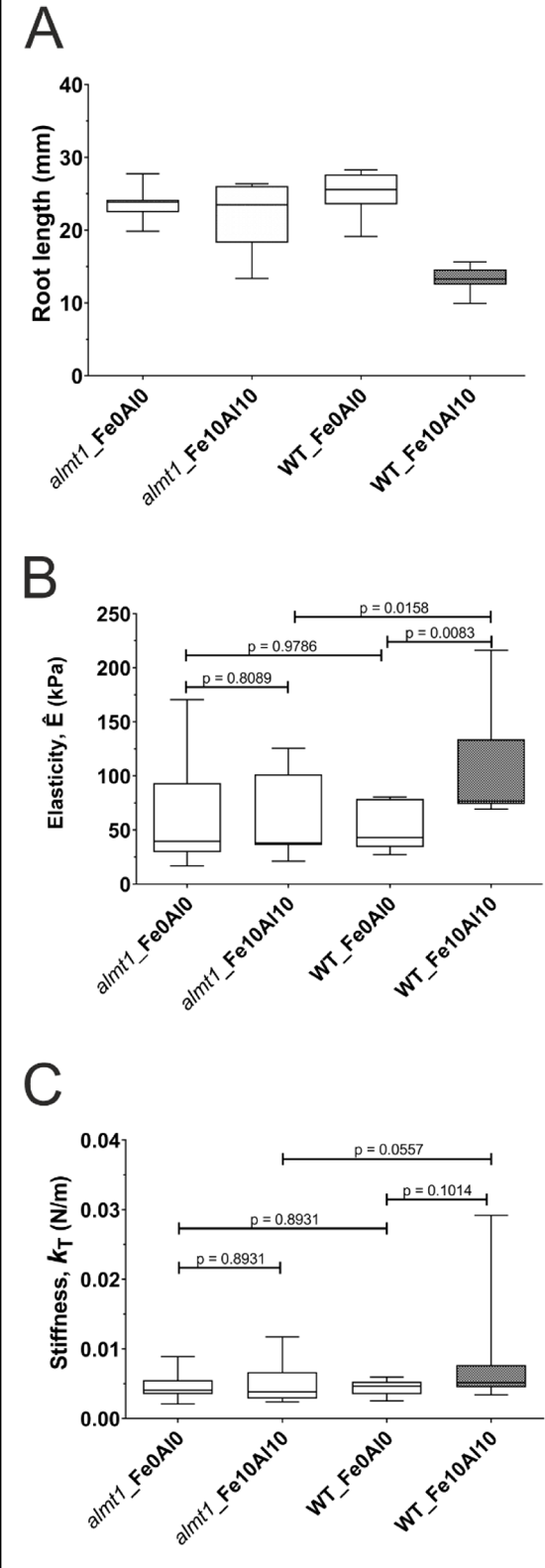
Box-and-whiskers plots of elastic properties of *almt1* mutant roots in comparison with WT (cf. Fig. 2) in two stressed conditions, Fe**0**Al**0** and Fe**10**Al**10. A**) Average root lengths measured on day 6. **B**) Effective Young’s modulus, *Ê*, (in kPa). **C**) Presentation of *k*T in the N/m unit

**Table 3:**
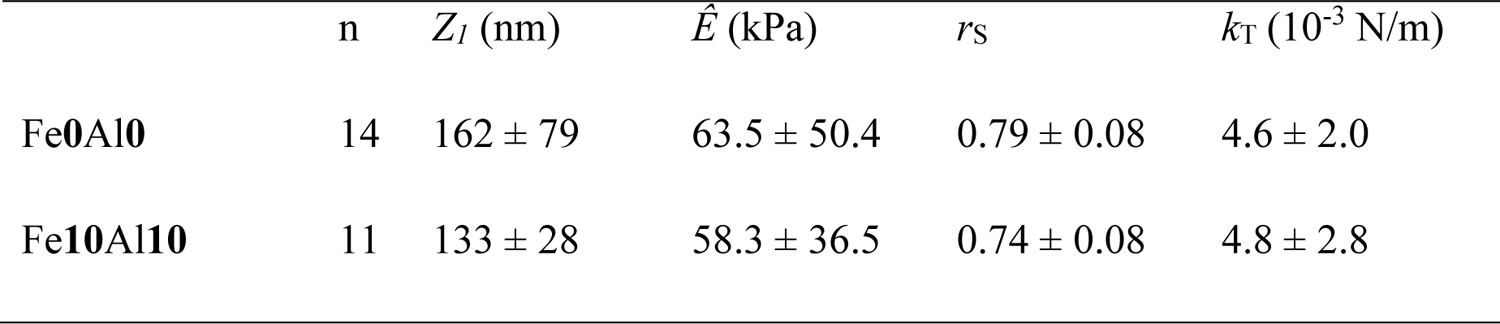
Elastic properties of *almt1* mutant seedling roots

| | n | $Z_l$ (nm) | $\hat{E}$ (kPa) | $r_s$ | $k_T$ ($10^{-3}$ N/m) |
| --- | --- | --- | --- | --- | --- |
| Fe0Al0 | 14 | $162 \pm 79$ | $63.5 \pm 50.4$ | $0.79 \pm 0.08$ | $4.6 \pm 2.0$ |
| Fe10Al10 | 11 | $133 \pm 28$ | $58.3 \pm 36.5$ | $0.74 \pm 0.08$ | $4.8 \pm 2.8$ |

## Discussion

The present results show that the elastic responses of external epidermal cell walls of *Arabidopsis* seedling roots to external forces vary in terms of concentration and composition of Fe and Al metal ions. It indicates that elasticity of plant cell CW is sensitive and can be used as to assess abiotic stresses on plant growth and stiffening. However, unexpectedly, the stiffening and the phenotype of seedling roots such as REA are not directly correlated.

### Root extension arrest (REA) and metallic stress

The root lengths of *Arabidopsis* seedlings were measured from the root tip to the cotyledon base (**Fig. S2**). Among all the stress conditions (Fe**10**Al**0**, Fe**0**Al**10**, Fe**20**Al**0**, Fe**0**Al**20**, and Fe**10**Al**10**), we observed the REA phenotype appeared only in the WT roots grown in Fe**10**Al**10** condition (**Table 1**). It is surprising that no REA was observed with doubled concentrations of single metal species (either Fe or Al). It reveals that the excess of single metal species did not urge the occurrence of REA. To ascertain that the REA phenotype is only due to the mixture of the two metal species, we carried out an experiment in a condition with the same metal ingredients and 500 µM phosphate (Pi). Phosphate is known for binding cations (Foy, Chaney, & White, 1978) but do not completely abolish the entry of metals into seedling roots (Balzergue et al., 2017). Results show no REA in the presence of Pi (**Table 2**, **Fig. S5**).

To further resolve the origin of REA occurrence, the WT results were compared with those of the *almt1* mutant. Lacking the malate-transporter ALMT1, the *almt1* mutant is strongly altered in exuding malate, a small organic anion known to chelate Fe^3+^ and Al^3+^. The root growth of *almt1* mutant was known to be insensitive to Fe^2+^ (under -Pi condition) and exhibited no REA phenotype (Balzergue et al., 2017; Mora-Macias et al., 2017). The absence of REA phenotype was explained as a consequence of reduced accumulation of iron in the apoplast due to a dramatic decrease of malate exudation. From our results, the mixed Fe and Al stress also lacks the ability to stimulate REA in the *almt1* mutants, and these mixed metal cations act like single metal ions of 10 μM in WT roots. In other words, without the malate exudation, the mixed Al and Fe are no longer growth inhibitors, leading to a normal growth. It further suggests that trapping metal ions by malate molecules is a key step to promote REA in WT. Taken all the data together, the factors to simulate REA include the amount of metal ions, the composition of metal species and the exudation of malate.

### Metallic stress and elasticity of living seedling roots

When the interlaced architecture of CWs is perturbed by metal ions, the bonding modes are accordingly justified; these changes can be reflected by altered elastic responses. It is noteworthy that the used AFM indenting tip has a small apex (∼10 nm radius), enabling us to sense structural strengths of beneath constituents in primary CWs such as cellulose microfibrils. Applying the trimechanic-3PCS framework to data analysis, the elasticity parameters defined therein helped us to differentiate elastic properties modulated by various stressed environments. The force decomposition of the theory unveils that the *FS*-deduced *Ê* is a sensitive parameter to varying metal contents in the growing medium, whereas the values of *k*T, representing the overall stiffness, are less distinguishable (**Table 2**). The change in penetration ease *r*S underlies the varying modes of nanomechanics and network bonding of CW architecture under different stresses (Chen et al., 2023). The *r*S parameter is provided only by the trimechanic-3PCS framework and cannot be accessed by the conventional methods (Hermanowicz, Sarna, Burda, & Gabrys, 2014). This *r*S parameter can also represent the deformability of the indented root.

According to comparable *r*S values of WT roots in Fe**0**Al**0**, Fe**10**Al**0** and Fe**0**Al**10** conditions, the bonding properties of CW structure are inferred alike. However, with higher concentration of metal ions (Fe**20**Al**0**, Fe**0**Al**20**, and Fe**10**Al**10**), an increase in *r*S is visible (*r*S > 0.8, **Table 2**). It follows that in all these conditions of high metal amount, the bonding properties of CW are differentiated from that of low metal amount. As already demonstrated, Al binds directly to negatively charged pectins of CWs and provokes a reduction in CW extensibility (Ma et al., 2004; Yang et al., 2016). In addition, expression profiling experiments suggested that pectins do bind with Fe (Hoehenwarter et al., 2016), which therefore, like Al, changes the bonding elasticity of the external primary cell wall. It is noteworthy that elastic parameters presented here are referred to the indentation depth of about 150 nm, which locates most likely the pectin constituents of CW (as opposed to cellulose microfibrils). Thus, the increased stiffness of CW for seedlings grown from Fe**20**Al**0** and Fe**0**Al**20** conditions likely involve the binding of Fe and Al to the pectin components of CW.

From the results of *Z1*, *Ê* or *k*T, the increase of the total amount of metal ions is closely related to CW stiffening. At 10 μM of either iron or aluminum, the elastic properties of WT roots are similar to that of the control system that contains no metal ions. At 20 μM (regardless of metal composition), the parameters *Ê*, *k*T and *r*S increase while *Z1* slightly reduces; see the results from Fe**10**Al**10**, Fe**20**Al**0** and Fe**0**Al**20** in **Table 2**. However, the Fe**20**Al**0** and Fe**0**Al**20** (single metal species) conditions exhibit no REA phenotype. It ensues that the increase of CW stiffness is not causal or not sufficient to trigger to REA, at least not in the operational conditions of our experiments.

### Root extension arrest and CW stiffening

REA phenotype induced by the Al stress is multifactorial and its mechanism remains largely unknown (Kochian et al., 2015). However, from our previous work and others, REA phenotype due to Fe stress is documented in its initial steps of Fe redox cycle that produces ROS in the CW and promotes peroxidase-dependent cell wall stiffening in the transition zone (Balzergue et al., 2017; Naumann et al., 2022). The major tolerance mechanism of Al toxicity is through the stimulation of the expression of *ALMT1* gene (Godon et al., 2019), a malate transporter (ALMT1 (Sasaki et al., 2004)). The rate-limiting step in this mechanism is the transport of organic acids rather than the cellular synthesis of these molecules (Ryan, Delhaize, & Jones, 2001). Indeed, Al^3+^ binding to the extracellular face of the ALMT1 channel opens the channel thereby stimulating the exudation of malate (Wang et al., 2022).

We have shown that the CW stiffness increases without REA at high Fe^2+^concentrations (≥ 20 µM) for seedling roots grown from an agar medium with poor phosphate and other metals, probably reflecting a lack of ROS production (**Fig. 5**). In the ferrous state, the Fe ion has multiple possible outcomes: adsorbed by the cell via its importing receptor, chelated with some organic acids in the CW, and oxidized to a ferric ion that may non-specifically bind to pectins of the CW. However, at 20 µM Fe^2+^, none of these outcomes are important enough to form the necessary redox condition for REA occurrence. At the same high concentration, Al^3+^ activates the exudation of malate that chelates Al to move it out of the root. The remaining Al^3+^ ions in the CW then bind to negatively-charged pectins, leading to an increase of stiffness though without REA occurrence (**Fig. 5**). The stress effect of co-presence of Fe and Al highlights the importance of malate accumulated in the apoplast. A current model postulates that, in combination with the apoplastic ferroxidase LPR1, malate-Fe^3+^ complexes trigger ROS in the apoplast (Naumann et al., 2022). Based on this model, our results show that Al^3+^ increases exudation of malate in the apoplast, thereby accumulating Fe in the apoplast followed by an accumulation of ROS to end up with a root extension arrest (**Fig. 5**).

**Figure 5:**
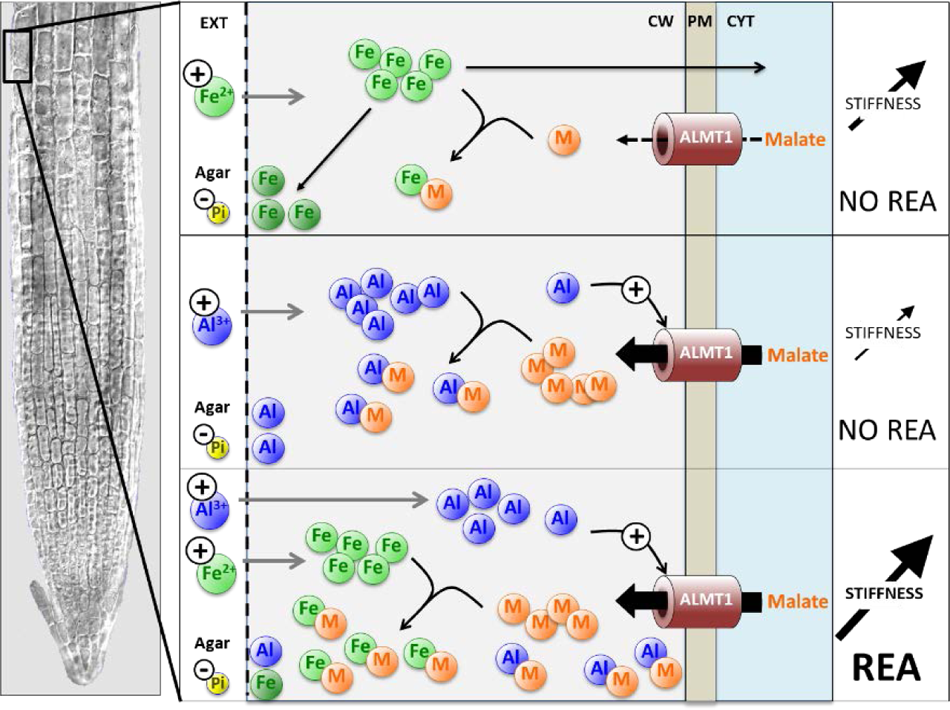
Model explaining the effects of Fe and Al on CW stiffening and root extension. Left panel shows a reconstituted picture of an Arabidopsis primary root tip; the square indicates part of the epidermis in the transition zone, where AFM measures were performed in this work. The top to bottom panels explain the phenomena that occur, depending on the Fe^2+^ and Al^3+^ content of the Pi-poor culture medium. Top panel: the Fe^2+^ ions enter the apoplast of the cell wall (CW, in light gray background color), which subsequently can cross the plasma membrane (PM, in light tan color) through an unknown transporter (not presented here for clarity) and activates the STOP1-ALMT1 signaling (not shown), or accumulate into the apoplast in complex with small organic acids like malate (M). The ALMT1 transporter exports malate from the cytosol (CYT, light blue background) to the CW. The accumulation of Fe cations, possibly in the Fe^3+^ state (darker green on the bottom left) binds to pectins, thereby increasing CW stiffness without triggering the root extension arrest (REA). Middle panel: the Al^3+^ ions enter the CW and activate the transcription of *ALMT1* (not shown) and the opening of ALMT1 transporter, thereby releasing malate in the apoplast. The accumulation of Al^3+^ leads to a modest increase of CW stiffness without REA. Bottom panel: the combination of Fe^2+^ and Al^3+^ results in a large release of malate and a high accumulation of ROS-promoting iron-malate complexes in the CW. These ROS concomitantly greatly increase CW stiffness and strongly prevents root extension. (M, malate; CW, cell wall; CYT, cytoplasm; PM, plasma membrane; REA, root extension arrest; -Pi, phosphate-poor medium; +Fe, adding Fe^2+^ in the medium; +Al, adding Al^3+^ in the medium)

## Conclusions

Root extension arrest was observed from *Arabidopsis* WT seedlings only stressed by a mix of 10 µM FeCl2 and 10 µM AlCl3 in a low phosphate agar medium. This REA is concomitant with a stiffening of the external primary cell walls. However, single metal, even at a higher concentration (20 µM), did not induce REA despite an increase in CW stiffness. Thus, the increase in the stiffness of CW may have independent origins: one associated with the binding of metals to pectin components of CW, and another associated with the redox cycle that produces ROS in the CW and promotes the peroxidase-dependent stiffening of CW. Consequently, the REA occurs in a balance of metabolic events (chemical and/or mechanical) that depends upon a change in the contribution of each factors including the chelating effect of malate in the combined Fe-Al stress.

## Acknowledgments

IBS acknowledges integration into the Interdisciplinary Research Institute of Grenoble (IRIG, CEA). This work acknowledges the AFM platform at the IBS. Acknowledgment to the ANR project BioPhyt-18-CE20-0023-03 and the support of the European Union’s Horizon 2020 research and innovation programme under the Marie Skłodowska-Curie grant agreement No 812772, Project Phys2BioMed. We thank Dr. Anne-Emmanuelle Foucher (IBS/EPIGEN) for critical improvements of the plant growth protocol. We also thank Isabel Ayala and Lionel Imbert (IBS/NMR) for their support in lab experiments.

## Author contributions

HK established the current protocol and performed the mechanical measurements. Jean-Marie Teulon performed the initial mechanical analysis. Christian Godon established the initial preliminary protocol and performed initial AFM measurements. Shu-wen W. Chen developed the trimechanic theory used to interpret the mechanical data. Thierry Desnos initiated the project and provided plant materials. Jean-Luc Pellequer directed the study and wrote the manuscript. All authors contributed to revisions of the manuscript.

## Data Availability

Data available on request from the authors

## Supplementary data of

### Preparation of agar

The nutrient solution contained 0.47 mM MgSO4, 2.1 mM NH4NO3, 1.89 mM KNO3, 0.67 mM CaCl2, 0.5 mM KI, 0.79 mM H3BO3, 10 mM MnSO4, 5 mM ZnSO4, 1 mM Na2MoO4, 0.1 mM CuSO4 and 0.1 mM CoCl2. The growth solution contains 20 ml/L of nutrient solution and 5 g/L sucrose. The agar medium contains 20 ml/L of nutrient solution with 5 g/L of sucrose and 8 g/L of agar powder (Sigma-Aldrich, A7921 Lot BCBZ7284). The elemental composition of the agar indicates a poor metal content (Mercier et al., 2021). The agar medium was buffered extemporaneously with 3.4 mM 2-(N-morpholino) ethanesulfonic acid (MES) for pH 5.5-5.8 range.

### Plant sealing under NuSil

A thin layer of silicone, NuSil MED1-1356 (NuSil Technology LLC, Carpinteria, CA, USA), was spread on the glass slide as described elsewhere (Kaur, Godon, Teulon, Desnos, & Pellequer, 2023). Partial polymerization was allowed for a few seconds before the root is laid over the silicone. Then, several thin silicone bands were stretched using a syringe needle to fasten the root over all its length except the transition zone, which is located about 500 µm from the root apex (**Fig. S1**). To prevent drying, a droplet of the growth medium (without the agar powder) was deposited to cover the entire seedling. After AFM calibration, the mounted seedling was positioned under the AFM for data acquisition (**Fig. S1**).

**Figure S1:**
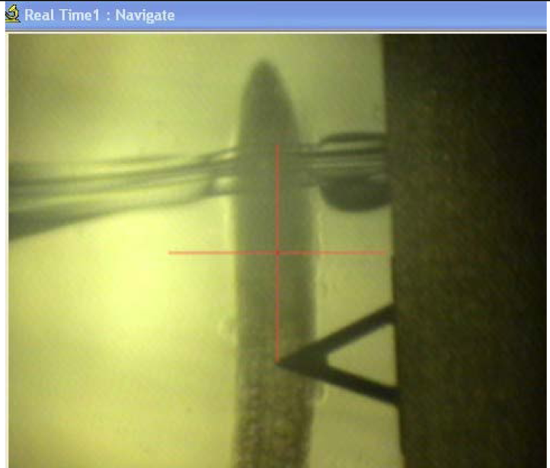
Photo of a root placed under the AFM cantilever taken by the AFM optical camera. The triangular shape cantilever (200 µm long) was placed 500 µm away from the root tip in the transition zone where nanoindentation measurements proceeded. The band of NuSil glue was near the root tip. The thickness of the fastening band must be thin enough to avoid hindering the AFM cantilever, but thick enough to withstand the bending of the root tip.

### Plant primary-root extension phenotype

Roots were transferred on day 4 in a Petri dish with stress agar medium, some of them were taken for nanoindentation experiments, while the remaining ones were labeled with a marker at their ends when the nanoindentation experiment finished. In the next day, a photo of the Petri dish is taken and archived with the experimental data. In **Fig. S2**, there is a clear demonstration of root extension arrest for the seedlings deposited in the agar medium with both Fe and Al, whereas the plant grew normally in the absence of metal stress.

**Figure S2:**
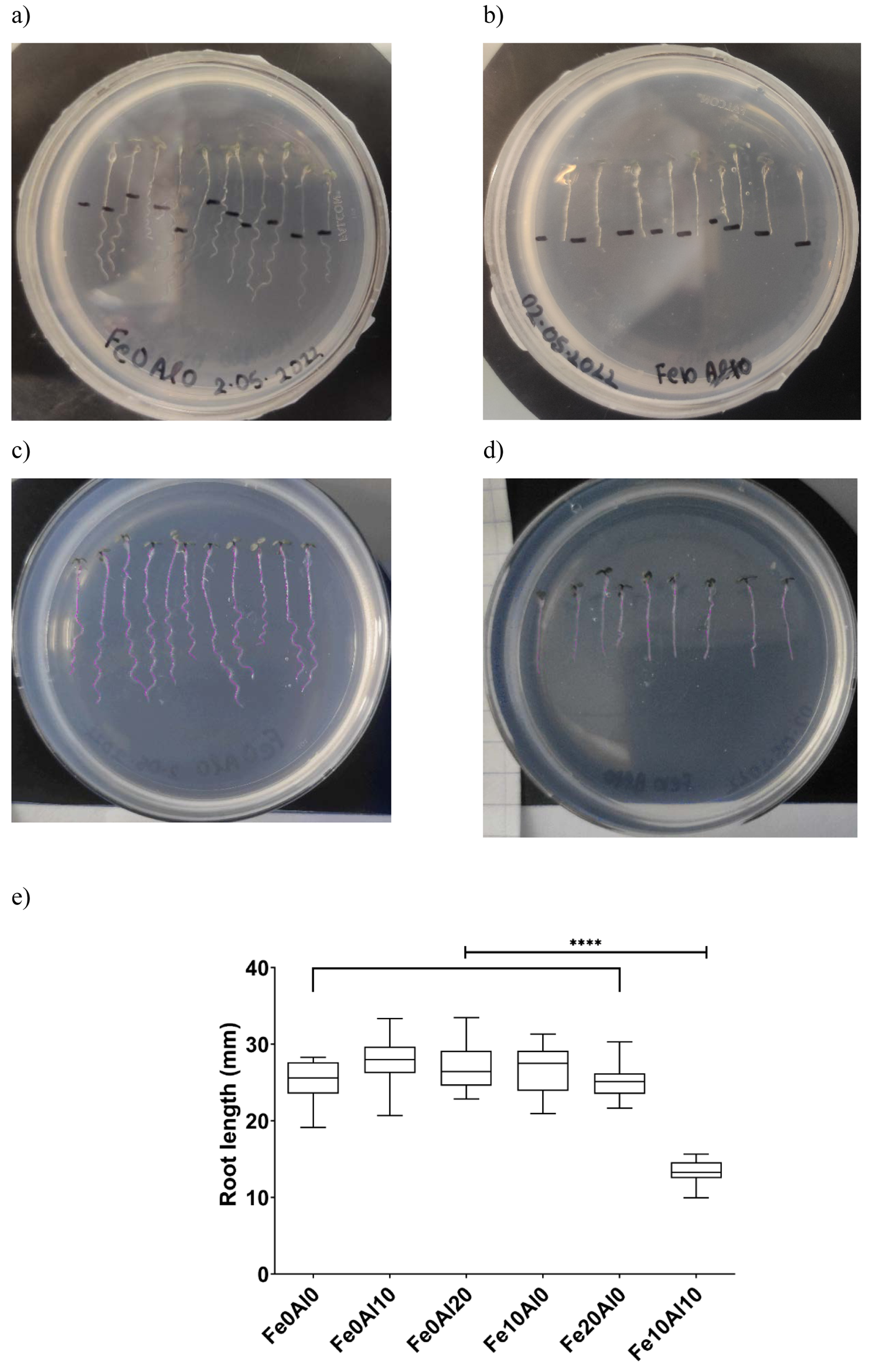
Root extension phenotypes. **a**) Control system: roots in the absence of metallic stress shows a normal root extension. **b**) Fe**10**Al**10** system: root extension arrest was observed by the lack of additional length measured below the marker. **c**-**d**) Snapshots for root length measurements using the NeuronJ plugin of ImageJ in the two systems of Fe**0**Al**0** and Fe**10**Al**10. e**) Box-and-whiskers plot of root lengths measured from all stress systems using NeuronJ expressed in mm units. Each box edge represents the first and third quartiles around the median. The whiskers represent the min and max values for each group. The nomenclature follows the metal stress conditions such as Fe**0**Al**0** for 0 µM of Fe^2+^ and 0 µM of Al^3+^, and so others for different combinations of both metals. Among all the metal stress conditions, only Fe**10**Al**10** shows a total root extension arrest, observed two days after AFM indentation experiments, and the statistical significance is labeled with **** toward every single condition (p < 0.0001).

On day 6, another photo was taken and saved as a GIF image format, which is compatible with the NeuronJ plugin (Meijering et al., 2004) of ImageJ software (Schneider, Rasband, & Eliceiri, 2012). NeuronJ was used to trace the whole root body until the base of the cotyledon to obtain the length (in mm) by an internal ImageJ calibration. The average of root lengths in different experimental conditions is shown in **Fig. S2E**.

### Data analysis by AtomicJ

For comparison, force-distance curves were also analyzed using AtomicJ (Hermanowicz, Sarna, Burda, & Gabrys, 2014), a standard nanomechanical analysis software. Parameters are as follows: robust exhaustive contact estimator with the robust (LTA) fitting method based on the Sneddon model for pyramidal tips of 35° half-opening. The approach force curve was capped to 5 nN. We regularly observed curves showing a poor R² fitting; we removed all the fitting data with R² < 0.9 (**Fig. S3**). Elasticity parameters for each measured node were grouped into a single value, each node produces an arithmetic average over all its force curves (4 or 9 curves), and finally all the arithmetic mean values from nodes were geometrically averaged to get one value per plant. Then, the final averages over all the plants from individual experimental conditions at one time were calculated and clubbed together (**Fig. S4**) and **Table S1**.

**Figure S3:**
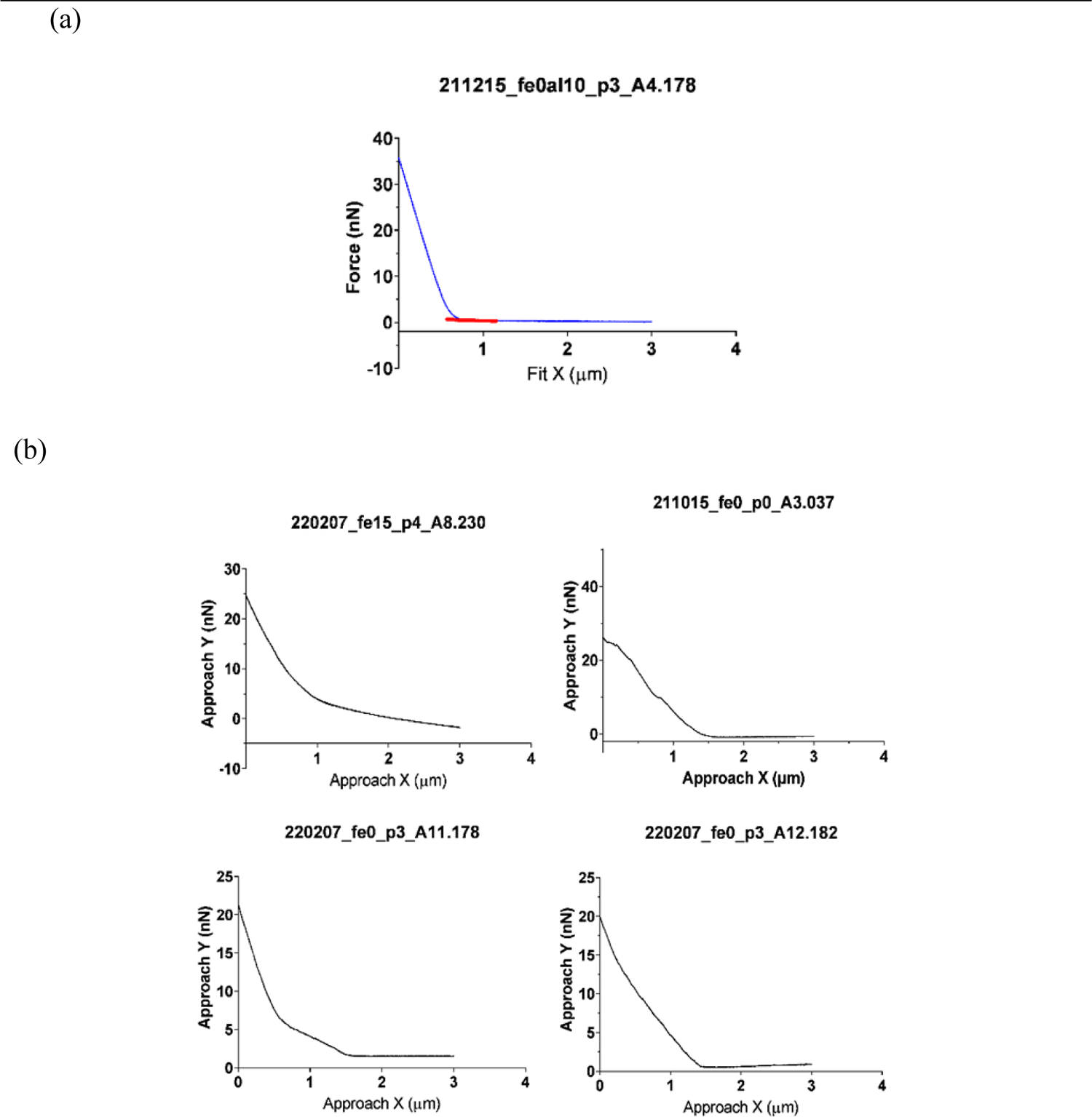
Representative force-distance curves excluded from AtomicJ analysis. **a**) A typical fine curve but with a poor fitting. Without optimizing the contact point manually, the curve was classified as a series of unqualified data and removed based on the value of R². From experiences, manually adjusting the location of the contact point for better curve fitting lacks an objective criterion for the user, leading to an unreliable outcome. Roughly, 5-10 curves were manually deleted per plant. If the number of deleted curves exceeded 30, then all the measurements from the plant were totally rejected and considered as a global poor data acquisition. **b**) Examples of excluded curves according to visual inspection. These curves visibly do not have a good approaching trace, therein several slopes or tilted baselines are present, indicating nanoindentation measurements were not properly acquired by AFM.

**Figure S4:**
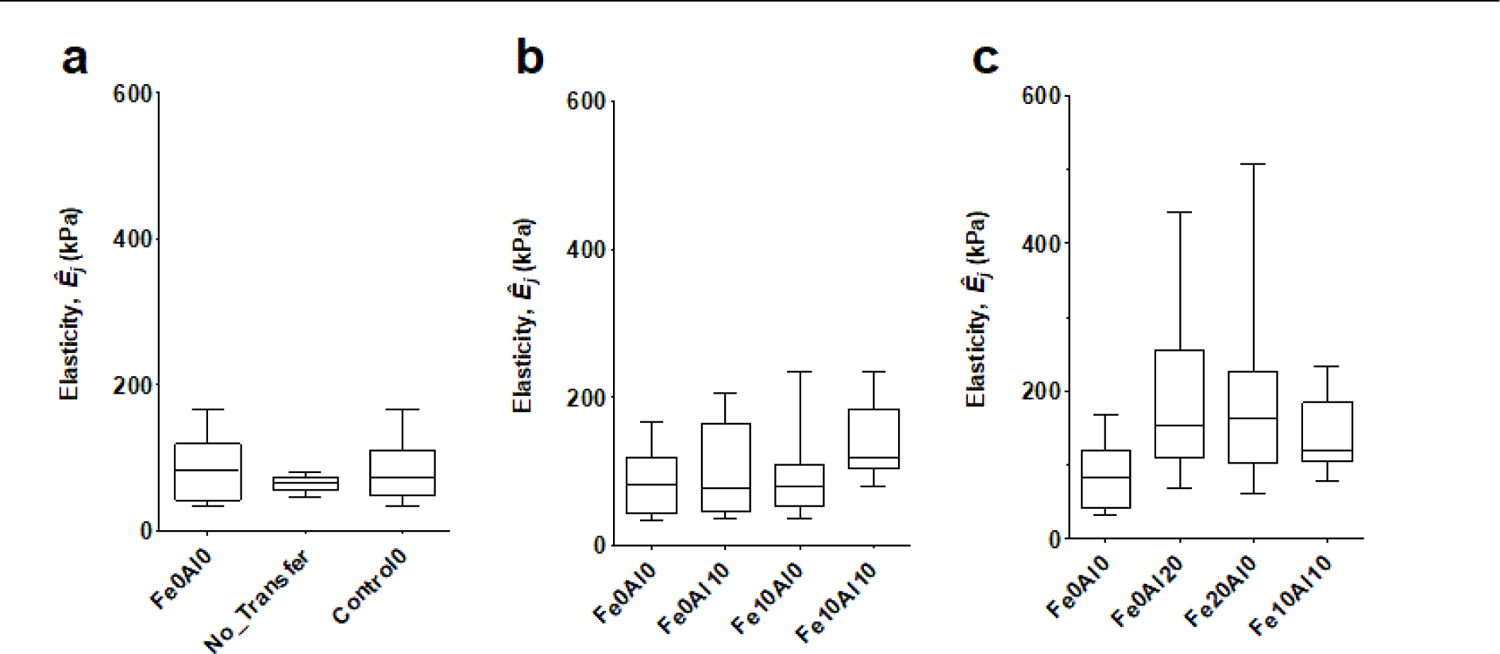
Box-and-whiskers plots of elastic properties of WT seedling roots analyzed by the pyramid model in the AtomicJ software. These plots **a-c** mirror those from **Figs. 1-3** in the main text. The effective Young’s modulus presented are in the kPa unit. See **Fig. S2** for numerical meanings of the box-and-whiskers plot. Fe**0**Al**0** indicates the nutrient solution without Fe and Al. No_Transfer denotes the seedlings lack the transfer step from crystallography plates to Petri dishes and elasticity nanoindentation experiments were performed directly after being taken out of the growing plates. Control0 is referred to the overall results from both Fe**0**Al**0** and No_Transfer conditions. In parallel, Fe**10**Al**10** contains 10 µM of Fe^2+^ and 10 µM of Al^3+^, and the rest of other stress systems can be perceived from their names.

Performing the statistical analysis using the same strategy, the elasticity results from the AtomicJ-pyramid method are globally similar to that from the trimechanic-3PCS framework. They show that under stress of Fe**10**Al**10**, the stiffness of CWs is higher than the other stress conditions. Moreover, the increase of metal amount (up to 20 µM) also stiffens the root. The data values from AtomicJ use a Poisson’s ratio of 0.5 which is different from the effective Young’s modulus used in the trimechanic-3PCS framework (*η* = 0); consequently *ÊJ* values from AtomicJ are, by definition, systematically 25% higher than *Ê* from the trimechanic-3PCS.

**Table S1:** The values of the effective Young’s modulus *ÊJ* by the AtomicJ-pyramid method

| Stress Conditions | $\hat{E}_J$ (in kPa) |
| --- | --- |
| Fe0Al0 | 87.7±45.3 |
| No_Transfer | 64.9±11.5 |
| Fe0Al10 | 99.9±65.9 |
| Fe10Al0 | 94.8±56.7 |
| Fe10Al10 | 141±57 |
| Fe20Al0 | 192±123 |
| Fe0Al20 | 186±112 |

### Nanomechanical measurement in presence of phosphate

Below are the nanomechanical results of WT seedlings when measured in presence of 500 µM of inorganic phosphate (Pi) using the trimechanic theory (**Fig. S5**). No significant change in elasticity, stiffness, or root length, are observed with Fe**10**Al**10** in presence of phosphate.

**Figure S5:**
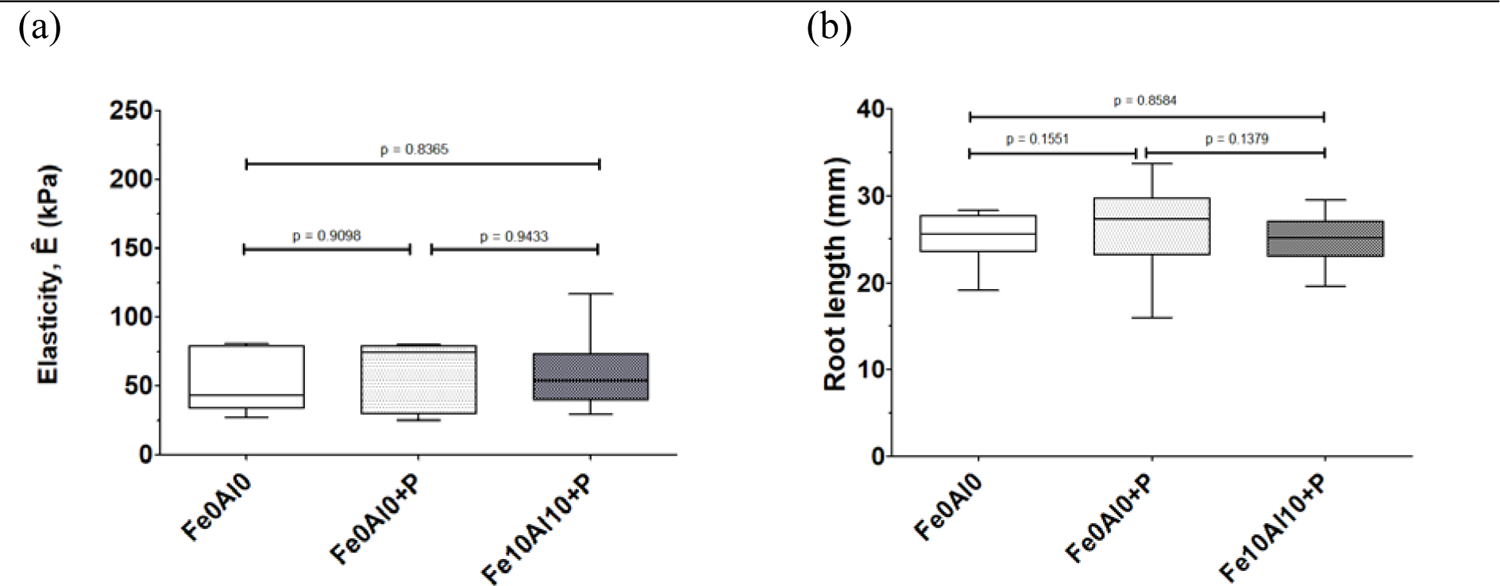
Box-and-whiskers plot of nanomechanical properties of WT seedling roots with the help of trimechanic theory. **a**) Elasticity values represented by the Young’s modulus expressed in kPa units. **b**) The lengths of seedling roots (in mm). See **Fig. S2** for the use of Box-and-Whiskers plots. The nomenclature of stress conditions follows the description presented elsewhere in the paper, where “P” in Fe**0**Al**0**+P denotes 500 µM of inorganic phosphate (Pi) added in. There were 5 and 8 plants for the conditions Fe**0**A**l0**+P and Fe**10**Al**10**+P, respectively?

